# The in-vivo microstructural profile of human hippocampal subfield CA1 and its relation to memory performance

**DOI:** 10.64898/2026.03.30.714764

**Authors:** Dayana Hayek, Joseph Höpker Fernandes, Niklas Vockert, Berta Garcia-Garcia, Hendrik Mattern, Niklas Behrenbruch, Larissa Fischer, Avinash Kalyani, Juliane Doehler, Dorothea Hämmerer, Yeo-Jin Yi, Stefanie Schreiber, Anne Maass, Esther Kuehn

## Abstract

The hippocampal CA1 subregion supports learning, memory formation, and spatial navigation. Although its three-layered architecture has been described in ex-vivo investigations, the in-vivo microstructural profile of CA1 and its relation to individual variations in memory performance remain poorly characterized. In this study, we used ultra-high field structural MRI at 7 Tesla to investigate the depth-dependent myelination patterns (measured by quantitative T1) of CA1 in younger adults, their relation to the local arterial architecture, and their association with individual differences in cognitive functions, specifically memory performance. Results show that left and right CA1 present depth-dependent patterns of myelination, with the outer and inner compartments showing higher myelination than the middle compartment. No significant relationship between layer-specific myelination of CA1 and distance to the nearest artery was observed. Right CA1 was found to be more myelinated than left CA1. Pairwise correlations and regression models showed that higher left CA1 myelination is linked to higher accuracy in object localization. Together, our data demonstrates the feasibility of describing the three layered myelin architecture of CA1 in vivo, and provides information on how alterations in the architecture of CA1 may relate to alterations in cognitive performance in younger adults.

## Introduction

The hippocampal CA1 region in the medial temporal lobe (MTL) is a pivotal component of human brain circuits, critically involved in learning, memory formation, and spatial navigation (Maass et al. 2014; Bein et al. 2020; Bobinski et al. 1998; Buzsáki and Moser 2013; Schapiro and Turk-Browne 2015; O’Keefe and Dostrovsky 1971). CA1 is known for its dense network of excitatory and inhibitory connections that are particularly susceptible to pathological changes in various neurodegenerative diseases (Xu et al. 2023; Bobinski et al. 1998), e.g., they are a main site of neuronal loss and gray matter atrophy in Alzheimer’s Disease (AD) (Duvernoy 2004; Yushkevich et al. 2024; Davies et al. 1992; de Flores et al. 2015), and lesions in this region relate to severe cognitive impairments, specifically anterograde amnesia (Zola-Morgan et al. 1986). CA1 is also an early site of tau accumulation (Braak and Braak 1991; Braak et al. 2006) and 7 Tesla in vivo and ex vivo MRI studies have shown that the apical layers (hippocampal stratum radiatum, lacunosum and moleculare; SRLM) are particularly vulnerable to AD-related atrophy (Kerchner et al. 2014, 2010; Boutet et al. 2014; Adler et al. 2018). However, relatively little is known about the in-vivo microstructural architecture of CA1 and its relation to individual memory performance. This is due to the difficulty of measuring small structures in the brain with 3 Tesla MRI technology.

Ex vivo investigations have shown that the microstructural profile of the CA1 region differs with respect to cell types, densities, and organization (Amunts et al. 2005; Insausti and Amaral 2004). In general, as the hippocampus is part of the archicortex, its cortical layers are classified into three broad cortical layer compartments compared to the six layers of the isocortex. The apical layers, including the SRLM, largely comprise the so-called plexiform layer receiving most inputs to the hippocampus. The somata of the principal pyramidal neurons form the middle pyramidal layer. The polymorphic layer, covering the basal layers like the stratum oriens and alveus, contains mostly axons, fibers, and interneurons that carry the output signals of the hippocampus (Insausti et al., 2017). Based on these cortical layer compartments, and given that the plexiform and polymorphic layers contain dense fiber tracts (such as the SRLM and alveus), which show greater myelination than the cell body–rich middle pyramidal layer, one may consider CA1 as a three-layered structure with respect to myelin distribution.

CA1 receives its main input from the apical layer III of the entorhinal cortex via the perforant path, terminating in apical dendrites located in the SRLM. Projections from CA2 and CA3, called Schaffer collaterals, terminate in stratum radiatum and stratum oriens. The major output of CA1 is sent from the basal layers (bordering the CSF) comprising the stratum pyramidale and stratum oriens (Insausti and Amaral 2004; Amaral and Lavenex 2006). The network architecture of CA1 is intricately linked to its microstructural properties, particularly the degree of myelination within its cortical layers, with the SRLM being highly myelinated (Kido et al. 1999). Myelination, the process of forming a myelin sheath around nerve fibers, is essential for the rapid and efficient transmission of electrical impulses along neural pathways, and may enhance the stability of functional representations (Kato et al. 2020; Das and Maharatna 2023). Myelin loss in cortical layers is a sign of pathology (Northall et al. 2024; Ward et al. 2014), and variations in myelination across different cortical layers can indicate aging and plasticity, and significantly impacts the overall performance of neural circuits and, consequently, sensory and cognitive functions (Sui et al. 2022; Northall et al. 2023; Liu et al. 2025).

Quantitative T1 (qT1) values (measured in milliseconds) can be used as a proxy for cortical myelination, and can be derived from MP2RAGE images (Marques et al. 2010). In regions that are high in myelin, qT1 times are shorter, therefore, qT1 is considered a myelin-sensitive measure (Stüber et al. 2014; Sprooten et al. 2019; Waehnert et al. 2016; Dinse et al. 2013). Whereas myelin is the main driver for the qT1 signal, iron explains around one third of the variance of cortical qT1 values (Stüber et al. 2014). By leveraging qT1 at 7T, we can obtain a quantitative, voxel-wise readout of microstructural variation in CA1.

Also 3T MRI studies have been critical in advancing our understanding of how MRI contrasts relate to biological tissue characteristics. Multi parameter mapping (MPM) demonstrates the feasibility of quickly acquiring robust and reproducible qMRI maps (longitudinal relaxation rate (R1), transverse relaxation ratios (R2*), Proton-Density (PD) and Magnetization Transfer Saturation (MTsat)) on 3T scanners at 1 mm isotropic resolution. The approach was further expanded to mitigate motion artifacts and improve acquisition efficiency (Weiskopf et al. 2013; Callaghan et al. 2016), and has revealed insights on microstructural cortex organization and aging (Callaghan et al. 2014; Karolis et al. 2019). qMRI metrics, such as R1, R2* and MTsat, have been investigated in 3T MRI studies (reaching submillimeter resolution of 0.8 mm), showing associations to cell-type specific gene expression patterns derived from spatial transcriptomic atlases (Wen et al. 2018; Patel et al. 2020; Edwards et al. 2023). Myelin water imaging (MWI) has been advanced to achieve high-quality myelin water maps at 2 mm isotropic resolution at 3T MRI (Shin et al. 2019). These studies demonstrate the potential of 3T MRI for analysing myelination in the cortex, including the MTL.

7 Tesla-based research has demonstrated that myelination levels can differ substantially between cortical layers, reflecting the expected pattern of depth-dependent myelination differences seen in ex vivo investigations (Doehler et al. 2023; Kuehn et al. 2017; Waehnert et al. 2016; Dinse et al. 2013; Liu et al. 2025). However, despite these advancements, there remains a gap in our knowledge regarding the specific distribution of myelination that can be detected in vivo in a small cortical area such as the hippocampal subfield CA1, and, prospectively, within other hippocampal subfields. Understanding the in vivo microstructural architecture of CA1, and its age- or disease-related alterations, will require both a more comprehensive characterization of CA1 architecture in the living human brain, as well as understanding the level of detail at which the microstructure of the circuit can be captured in vivo.

Consequently, we aimed to describe the depth-dependent myelination pattern of CA1 in vivo using a 7T MRI protocol that (i) can be measured in one scan, (ii) is optimized for structural hippocampal imaging, and (iii) adopts the methodology previously used for similar purposes in the neocortex (Doehler et al. 2023; Northall et al. 2023; Liu et al. 2025).

An architectural feature that has recently been shown to interact with local cortical microstructure is regional vascularization. Vascularization, and specifically the distance to the nearest blood vessel, may affect local tissue microstructure thereby influencing the qT1 signal. Myelin formation and maintenance impose high metabolic demands, and adequate vascular supply is critical for sustaining oligodendrocyte functions that are dynamic on a day-to-day basis (Philips and Rothstein 2017). Myelin sheaths close to arteries are expected to benefit more from regular metabolic supply, which may impact their architecture; in addition, myelin sheets may preferentially form in areas of high metabolic supply.

Prior work has shown that a dual hippocampal vascular supply pattern (i.e., a supply by two arteries) is associated with better cognitive performance compared to a single-supply pattern in patients with cerebral small vessel disease (Perosa et al. 2020), and that this relates to differences in hippocampal gray matter integrity (Vockert et al. 2021). Additionally, in the hippocampus, a link between vessel distance and local myelination has been shown, because higher distance to the supplying vasculature was related to lower myelination (Haast et al. 2023). Also in the primary motor cortex, the primary somatosensory cortex, and the prefrontal cortex, a 7T-MRI study has indicated a dependence of the local myeloarchitecture to the distance to the nearest blood vessel (Knoll et al. 2024). In order to understand if the distance to the nearest artery is a factor that explains variance in the (layer-specific) structural architecture of CA1, we included vascular distance as a factor in our model, in addition to age and sex.

Finally, we aimed to test whether a relationship exists between individual variation in layer-specific CA1 microstructure and individual cognitive performance. A link between different measures of hippocampal microstructure and cognition has been investigated by previous studies. In participants with early AD and mild cognitive impairment (MCI), microstructural alterations in the hippocampus, measured via mean diffusivity, significantly correlated with cognitive performance (Yakushev et al. 2011; Fellgiebel and Yakushev 2011). In a healthy cohort, however, no significant association between delayed recall performance and quantitative 3T MRI measures of microstructure (R1, MTsat, PD, R2*) were previously detected (Wearn et al. 2025). Although individual variations in the microstructure of CA1 are expected to be smaller in younger compared to older adults, individual differences that can be depicted with 7T MRI could explain variations in memory performance, providing critical indications for future points of vulnerability or as targets for interventions.

Taken together, we here offer an in vivo characterization of the microstructural profile of human CA1 in younger, healthy adults using 7T-MRI in combination with behavioral memory tests, aiming to explore the potential of 7T-MRI to detect layer-specific myelin differences in this region in vivo and their relation to behavior. We further aim to understand the contribution of specific variables on this architecture, specifically age, sex, hemisphere, and distance to the nearest blood vessel. Our data may therefore aid future studies to investigate how factors that are expected to influence the microstructural architecture of CA1, such as aging and neurodegenerative diseases, affect the layer-specific myelination of CA1 in the living human brain.

## Methods

### Participants

N=33 younger adults were initially included in the study. n=3 participants had to be excluded due to motion artifacts in the ToF images. The finally analyzed sample included n=30 younger adults (age: M = 26.04 years, SD = 3.72 years, ranging from 20 years to 36.5 years, 16 females). Participants were recruited from the participant database of the German Center for Neurodegenerative Diseases (DZNE), Magdeburg, Germany. Exclusion criteria for participation were 7T-MRI contraindications (e.g., presence of active implants, non-removable metallic objects, tinnitus or hearing impairments, consumption of alcohol or drugs before the measurement), and chronic illness. Participants underwent three appointments in total: (1) An MRI scanning session, where also cognitive data was acquired during the scan, (2) a behavioral cognitive testing session one week later, and (3) a second MRI scanning session 0.25-2.5 years later, which included the structural scans used in this study. The larger gap between (2) and (3) is due to the COVID-19 pandemic during which no MRI sessions took place for safety reasons. All participants were paid for their attendance and provided written informed consent. The study was approved by the Ethics committee of the Medical Faculty of the University of Magdeburg (193/19).

### Cognitive Testing

All participants underwent cognitive testing. During the first MRI session (1), participants performed a visuospatial memory task for object location (object location task (OLT)). First, participants completed an encoding session on a desktop computer. About one hour later, the first retrieval session was performed inside the fMRI scanner. One week later, during the behavioral session (2), participants underwent a cognitive test battery, composed of 7 tests: the Rey-Osterrieth Complex Figure Test (ROCFT) (Meyers and Meyers 2007), the Trail Making Test (TMT) (Reynolds 2002), the Logical Memory from the Wechsler Memory Scale (WMS) (Wechsler 1987), the Digit Span (DS) from the Wechsler Adult Intelligence Scale (Wechsler 2008), the Color and Word Interference Test after Stroop (SCWT) (Stroop 1935), and the Symbol Digits Modalities Test (SDMT) (Smith 2002). They performed the second retrieval session of the OLT on a computer screen. Scoring for the ROCFT was based on (Lezak 2004). For the recognition section of the ROCFT, a corrected hit rate was calculated by subtracting false positives from hits and then dividing by the number of true positives.

A composite memory score was calculated as the sum of z scores of the five memory tests: *f mem z = scale(OLT DR1) + scale(OLT DR2) + scale(ROCFT IR) + scale(ROCFT DR) + 2*scale(LM DR tot) (DR = Delayed Recall, IR = Immediate Recall, tot = Total).* The delayed recall from logical memory (LM DR) was included twice to give more weight to the verbal memory component, which would otherwise be underrepresented. For simplicity, we will henceforth refer to the composite memory score *f mem z* simply as *Memory*.

### MRI Image Acquisition

7T MRI data were acquired at a Siemens MAGNETOM scanner located in Magdeburg, Germany, using a 32-channel head coil. MP2RAGE whole-brain images were acquired at 0.5 mm isotropic resolution (352 sagittal slices, field-of-view read = 224 mm, repetition time = 4800 ms, echo time = 2.18 ms, inversion time TI1/TI2 = 900/2750 ms, flip angle = 5**°**/3**°**, bandwidth = 250 Hz/Px, GRAPPA 2, ca. 17 min).

7T T2-weighted images (optimized for MTL volumetry) were acquired at a 0.4375 x 0.4375 x 1.1 mm anisotropic resolution (50 coronal slices orthogonal to the hippocampal long axis, field-of-view read = 224 mm, repetition time = 8000 ms, echo time = 92 ms, flip angle = 60**°**, bandwidth = 158 Hz/Px, ca. 8 min). 7T Time-of-Flight (ToF) angiographic images were acquired at a resolution of 0.28 mm isotropic (176 transversal slices, field-of-view read = 220 mm, repetition time = 22 ms, echo time = 4.59 ms, flip angle = 5**°**, bandwidth = 142 Hz/Px, GRAPPA 3, slice oversampling = 18.2%, ca. 14 min). To mitigate the increased specific absorption rate at 7T MRI, sparse venous saturation was implemented following the approach of Schmitter (Mattern et al. 2018). More sequences were included in the protocol but were not relevant in this study (see Supplemental **Figure 1** for details). Scanning time summed up to around 70 minutes.

All scans underwent visual quality control and were inspected for artifacts, including image distortions and motion artifacts. In n=3 ToF images, motion artifacts were detected. Those participants were excluded from the study.

### Hippocampal Subfield Segmentation and Co-registration

A segmentation of hippocampal subfields was computed using the Automated Segmentation of Hippocampal Subfields (ASHS) software using the 7T Atlas for T2-weighted MRI of younger adults as input (Berron et al. 2017; Yushkevich et al. 2015). This atlas facilitates the segmentation of hippocampal subfields (such as subiculum, dentate gyrus, Cornu Ammonis 1, 2, and 3) and parahippocampal regions (including the entorhinal cortex, Brodmann areas 35 and 36, and parahippocampal cortex), aligning with the manual segmentation protocol established by Berron et al. (Berron et al. 2017). The automated segmentations underwent visual quality control. Manual curation (by N.V. and D.H.) was conducted, specifically targeting correction of three commonly identified errors (Xie et al. 2019): (i) under-segmentation of the lateral hippocampal border, (ii) misidentification of the choroid plexus as part of the hippocampus, and (iii) excessive segmentation of the medial temporal lobe cortices. Additionally, it was ensured that the borders of the segmentation do not extend beyond CA1, into adjacent white matter regions.

Both the hippocampal head and tail were excluded from the segmentation. These delineations are considered anatomically less reliable because key internal landmarks are less consistently visible and morphological variability is greater in the anterior portion (Wisse et al. 2016; Yushkevich et al. 2015). Moreover, the hippocampal tail is not segmented into individual subfields in standard ASHS protocols but treated as a single label (Yushkevich et al. 2015).

The whole-brain qT1 image computed from the MP2RAGE (https://github.com/JosePMarques/MP2RAGE-related-scripts) was registered to the slab T2 image using the ITK-SNAP (Yushkevich et al. 2006) manual registration tool. The registration involved an affine transformation with 6 degrees of freedom. Manual registration was chosen in order to ensure precise spatial alignment that could not be ensured using a fully automated pipeline.

### CA1 Depth Segmentation (Layer Analyses Pipeline)

Accurate and precise tissue segmentation is a crucial step to investigate CA1 layers. Note that the term “layers” as used here should not be interpreted as histologically-defined layers; instead, the term is employed to refer to 3 different compartments that we expect to align with expected myelination patterns (from apical, covering the SRLM, to basal, covering the basal pyramidal layers and the stratum oriens, followingly referred to as inner - outer, see Supplemental **Figure 2**). Layer analyses were performed using the LAYNII toolbox (version 2.9.0, Huber et al. 2021). The binary CA1 masks were upsampled to 0.2 mm isotropic image resolution. qT1 images were then cropped to the size of the bounding box of subfield CA1, and upsampled to the same image resolution using nibabel’s (Brett et al. 2024) resample_from_to() function with nearest-neighbour interpolation. This step allowed a smoothed calculation of CA1 features (like curvature and thickness).

The preprocessed qT1 images were analyzed using LAYNII to extract depth-dependent data points from the CA1 subfield. Initially, the CA1 subfield was divided into 21 equidistant depths using the ‘LN2_LAYERS’ command, facilitating a detailed depth-dependent structural analysis. Subsequently, the 21 depths were aggregated into three broader layer compartments, by merging the seven most inner depths (closest to DG) into an inner compartment, the seven most basal depths into an outer compartment (closest to CSF) and the remaining 7 depths into a middle compartment. See **Figure 1** for a summary of the analyses pipeline.^1^

**Figure 1.**
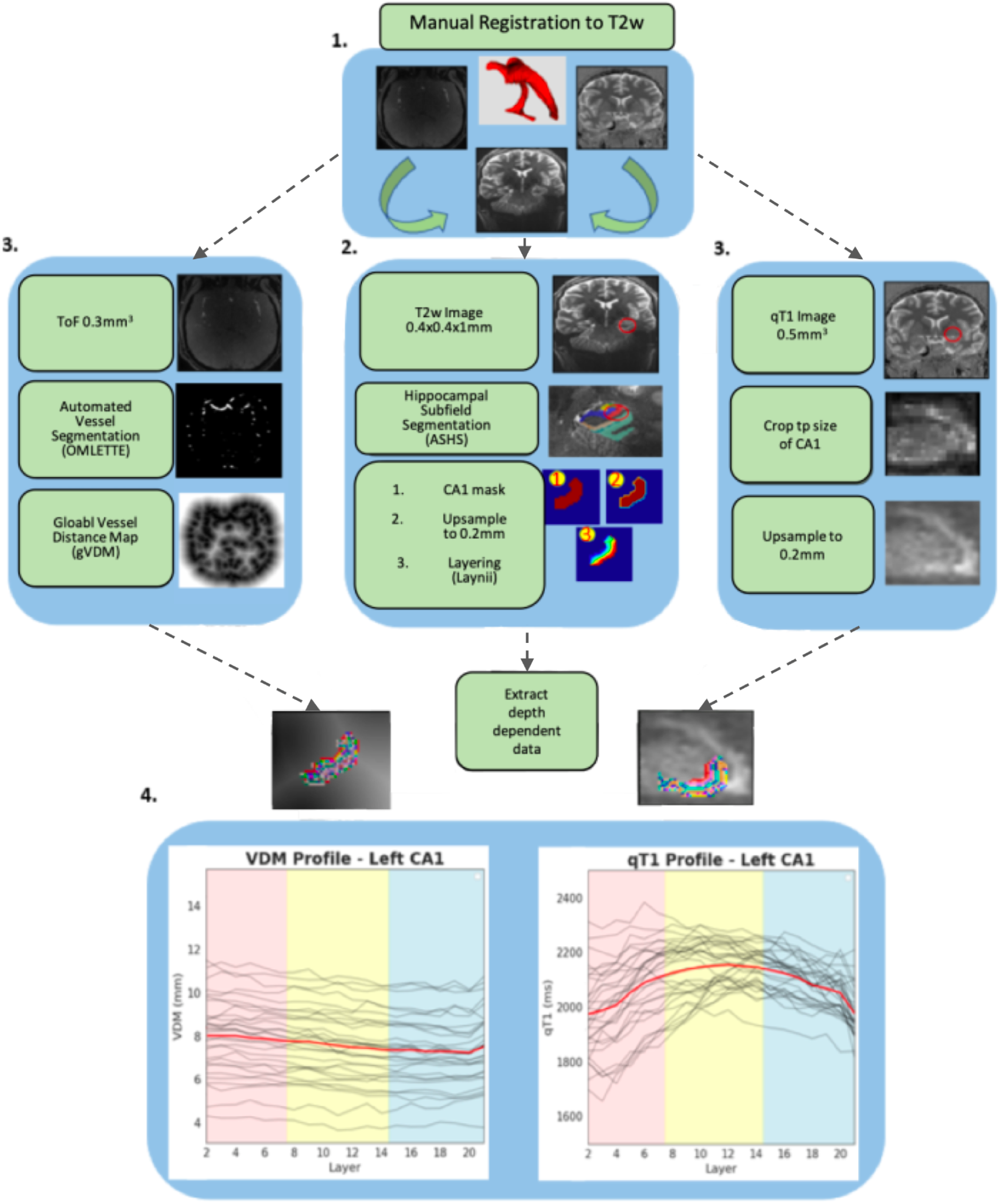
Hippocampal Depth Segmentation Pipeline. **(1)** qT1 whole-brain and Tof slab images were co-registered to the T2-weighted image using ITKsnap (Yushkevich et al. 2006)**. (2)** Automated Segmentation of Hippocampal Subfields (ASHS) was used to segment the hippocampus. The binary mask of the CA1 subfield was extracted and up-sampled. Then the inner and outer rim was defined on the mask, after which the rimmed mask was used to generate 21 equidistant layers using Laynii. This and the following steps are only shown for the left CA1 here. **(3)** Automated vessel segmentations were generated from the ToF images, using the OMLETTE toolbox. From the Vessel segmentations, global vessel distance maps (g-vdm’s) were computed. The g-vdms were cropped and resampled to the size and resolution of the CA1 layer file. Also, the qT1 image was cropped to the size of the bounding box of the CA1 subfield, and the crop was upsampled to match the resolution of the CA1 layer file. **(4)** Layer Dependent qT1 and VDM values were extracted.

### Vessel Segmentation and Vessel Distance Mapping (VDM)

To extract the vessels from the ToF images, we used the Openly available sMall vEsseL sEgmenTaTion pipelinE (OMELETTE, (Mattern 2021), https://gitlab.com/hmattern/omelette). After top-hat transformation, the vasculature was segmented using the Jerman filter given superior segmentation performance compared to other vessel enhancement filters, particularly for small vessels (Jerman et al. 2015). The parameter tau was set to 0.5 based on empirical optimization. Resulting vessel probability maps (representing the probability of each voxel belonging to a vessel) were thresholded using a 3-class Otsu analysis. Voxels above the upper threshold were classified as vessels, voxels below the lower threshold were classified as background. Hysteresis thresholding was employed to classify vessels between the lower and upper thresholds (Fraz et al. 2011), where a voxel is considered to belong to a vessel only if it is connected to a voxel with a value exceeding the upper threshold.

To compute the VDMs as described by (Garcia-Garcia et al. 2023), individual vessel probability maps were converted to binary format. Then, for the VDMs, the euclidean distance of each voxel to the closest segmented vessel voxel (in the binary vessel map) was calculated (for a visualization, see Supplemental **Figure 3**). To compute the distribution of vessel radii, skeletonization was performed on the binary vessel masks to determine the center voxels of the vessel. Then, an euclidean distance transform was used to determine the distance to vessel borders. The resulting histogram can be seen in Supplemental **Figure 4**.

### Statistical Analyses

Statistical analyses were performed using python and the statsmodels library (version 0.14.4, (Seabold and Perktold 2010)). A linear mixed model (LMM) was computed to assess the differences in myelination across three distinct layer compartments of the left and right CA1, including fixed effects for age and hemisphere, as well as compartment*hemisphere interactions. The model included random intercepts for subjects to account for within-subject correlations (qT1 ∼ Compartment * Hemisphere + age + sex + (1 + Compartment | Subject)). To investigate the effect of arterial distance on myelination, a LMM was used with qT1 values as the dependent variable and random intercepts to account for repeated measures. The analysis includes distance to the nearest artery (VDM) as a continuous variable, ranging from 0 - 14 mm as within-subjects factor with fixed effects for sex and age and hemisphere, testing also for VDM*compartment interactions (qT1 ∼ VDM * Compartment + Hemisphere + age + sex + (1 + Compartment | Subject)) and VDM * hemisphere interactions (qT1 ∼ VDM * Hemisphere + age + sex + (1 + Hemisphere | Subject)) in separate models. Pairwise correlations were computed to investigate possible relationships between depth-dependent CA1 qT1 values; resulting p-values were FDR corrected. Subsequently, a linear mixed-effects (LME) model was used to assess whether these relationships remained significant when accounting for covariates and random effects, also testing for myelination * hemisphere interactions (qT1 ∼ Memory * Hemisphere + VDM + age + sex + (1 + | Subject)). To further examine hemispheric effects, additional models were used to test for the effect of Memory on qT1 in the left and right CA1 separately (qT1 ∼ Memory + VDM + Age + Sex + (1+ | Subject), as well as testing for interactions with layer compartment (qT1 ∼ Memory * Compartment + VDM + Age + Sex + (1+ | Subject)). The significance threshold was set to p < 0.05.

## Results

### Myelination differences between CA1 depths and hemispheres

A linear mixed model was computed to examine the differences in qT1 values (higher values reflecting lower myelination) across three layer compartments of the right and left CA1, adjusting for age and sex. The model includes fixed effects for layer, hemisphere, age, and sex, and random intercepts for subjects to account for within-subject variability. A likelihood ratio test comparing models with and without hemisphere revealed significant main effects of hemisphere on qT1 values *(LRT = 159.57, df = 3, p < 0.0001, left hemisphere: mean qT1 = 2088.1 ms, std = 118.5, right hemisphere: mean qT1 = 1859.15 ms, std = 98.6)*, indicating higher myelination of CA1 in the right compared to the left hemisphere. There were no statistically significant interactions between layer compartment and hemisphere (middle × right: *β = -31.57 [95% CI -102.15, 39.01], SE = 36.01, p = 0.381*; outer × right: *β = 24.9 [95% CI -45.67, 95.47], SE = 36, p = 0.49*), suggesting that the relationship between hemisphere and qT1 values does not vary significantly across layer compartments. To probe if field inhomogeneities during the MRI acquisition contribute to the observed asymmetry, we compared average qT1 values between the left and right hemispheres of a control region. As a control region, BA35 was selected. The results of this analysis can be seen in Supplemental **Figure 5**. Additional analyses on a control region, BA35, show that there is a consistent hemispheric difference in qT1 values both in CA1 and BA35, with the right hemisphere showing lower average qT1 than the left (*CA1_left: Mean = 2102.4ms, CA1_right: Mean= 1876.3ms, %Δ_left-right_ = 10.6; BA35_left: Mean = 2010.5ms, BA35_right = 1778.1ms, %Δ_left-right_ = 11.6*). Age and sex were not significant predictors in the model (*sex: β = -2.75, p = 0.94; age: β = 1.48, p = 0.73).* Random intercepts show moderate subject-level variability (ICC = 0.307) Post-hoc comparisons reveal that qT1 values are significantly higher (indicating lower myelination) in the middle layer compartment compared to both the outer layer compartment (*middle - outer: t = 4.99, p < 0.001*) and the inner layer compartment (*inner - middle: t= -4.60, p < 0.001*) across hemispheres. A consistent laminar difference was hence detected, with the outer layer compartment showing the greatest myelination, followed by the inner and the middle compartments (see **Figure 3**).

**Figure 3.**
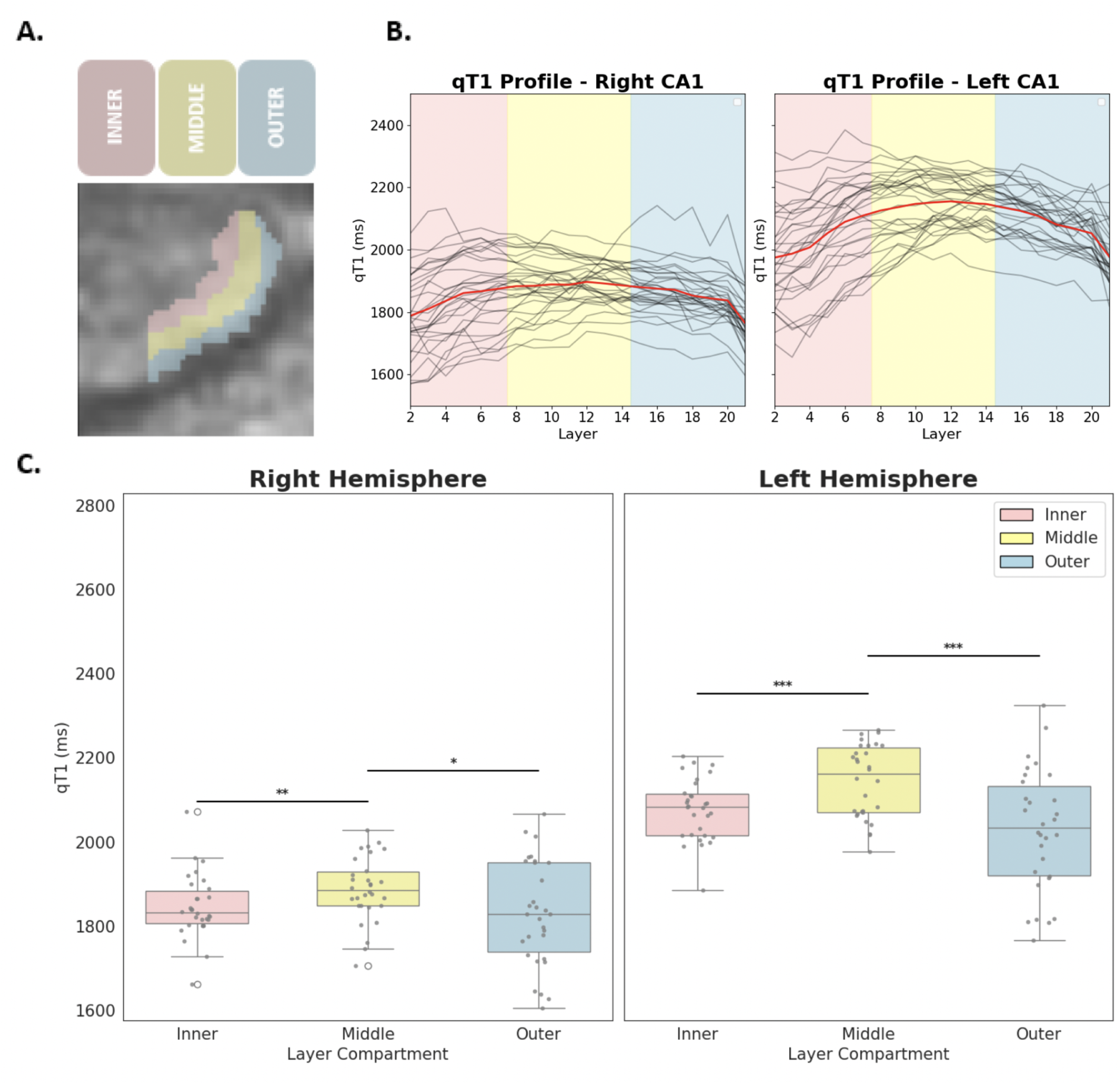
Relationship between CA1 depth and myelination. **A)** Positions of layer compartments on CA1(left): Blue: outer compartment (layers 1-7) close to CSF. Yellow: middle layer compartment (8-14). Red: inner layer compartment (layers 15-21) close to DG. **B)** Layer-specific myelination in human CA1. Layer myelin profile of all participants, in 21 depths of the left and right CA1. For every single layer compartment, the mean T1 time of all the voxels in that compartment is plotted. Lower T1 times indicate higher myelination levels. **C)** Boxplots show the distribution of myelin in each compartment for both hemispheres. Asterisks indicate significant difference between compartment mean compartment myelination (*** p<0.001, ** p<0.01, * p<0.05).

### No significant effect of vessel distance on CA1 myelination

However, the linear mixed-effects model reveals no significant effect of vessel distance on qT1 values (*β = -2.08 [95% CI -18.165, 14], STE = 8.02, p = 0.8*). In addition, there are no significant interactions between VDM and layer compartment (VDM × middle: *β = -0.918 [95% CI -21.23, 19.39], SE = 10.36, p = 0.93*; VDM × outer: *β = -15.518 [95% CI -35.75, 4.71], SE = 10.32, p = 0.133*), suggesting that qT1 values in different depths do not systematically vary in relation to their distance to the nearest artery. Also hemisphere does not interact with the relationship between VDM and qT1 values (*β = 3.2 [95% CI -17.9, 24.31], SE = 10.77, p = 0.76)*. Age and sex are significant predictors neither (age: *β =* 0.82, *p* = 0.19; sex: *β = -2.8 p* = 0.93, see **Figure 4**). Overall, vessel distance shows no significant effect on depth-dependent qT1 values in CA1 in our sample (composed of younger, healthy adults).

**Figure 4.**
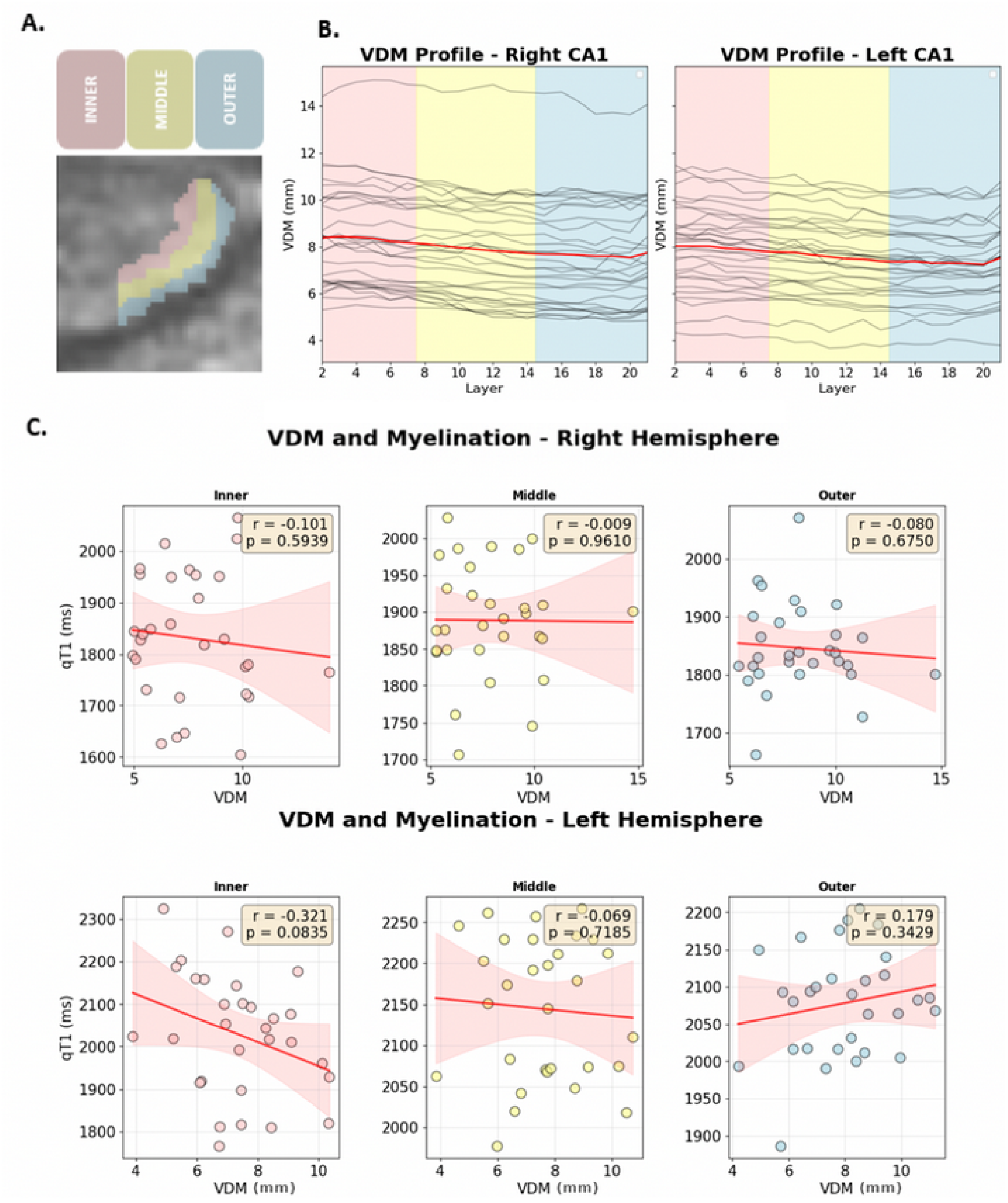
Relationship between Vessel Distance and Compartment Myelination. **A)** Positions of layer compartments on CA1(left): Blue: outer compartment (layers 1-7) close to CSF. Yellow: middle layer compartment (8-14). Red: inner layer compartment (layers 15-21) close to DG. **B)** Layer-specific vessel distance in human CA1. **C)** Relationship between myelination and vessel distance in each compartment. 95% Confidence Intervals indicated in Red. Pearson’s r- and p-indicated in yellow.

### Relationship between memory scores and CA1 myelination

To test for a potential relationship between individual differences in layer-specific myelination and memory performance, pairwise correlations were computed. After FDR-correction, they revealed significant negative correlations between the composite memory score and qT1 values in all three layer compartments of the left hemisphere (inner × Memory: *r_FDR_ = -0.35, p = 0.001,* middle × Memory: *r_FDR_ = -0.45, p < 0.001,* outer × Memory: *r_FDR_ = -0.26, p = 0.02,* see **Figure 5B**), indicating that better memory scores relate to higher myelination in all layer compartments of left CA1. For right CA1, no significant correlations were detected (also not significant without FDR-correction, see **Figure 5A**).

**Figure 5.**
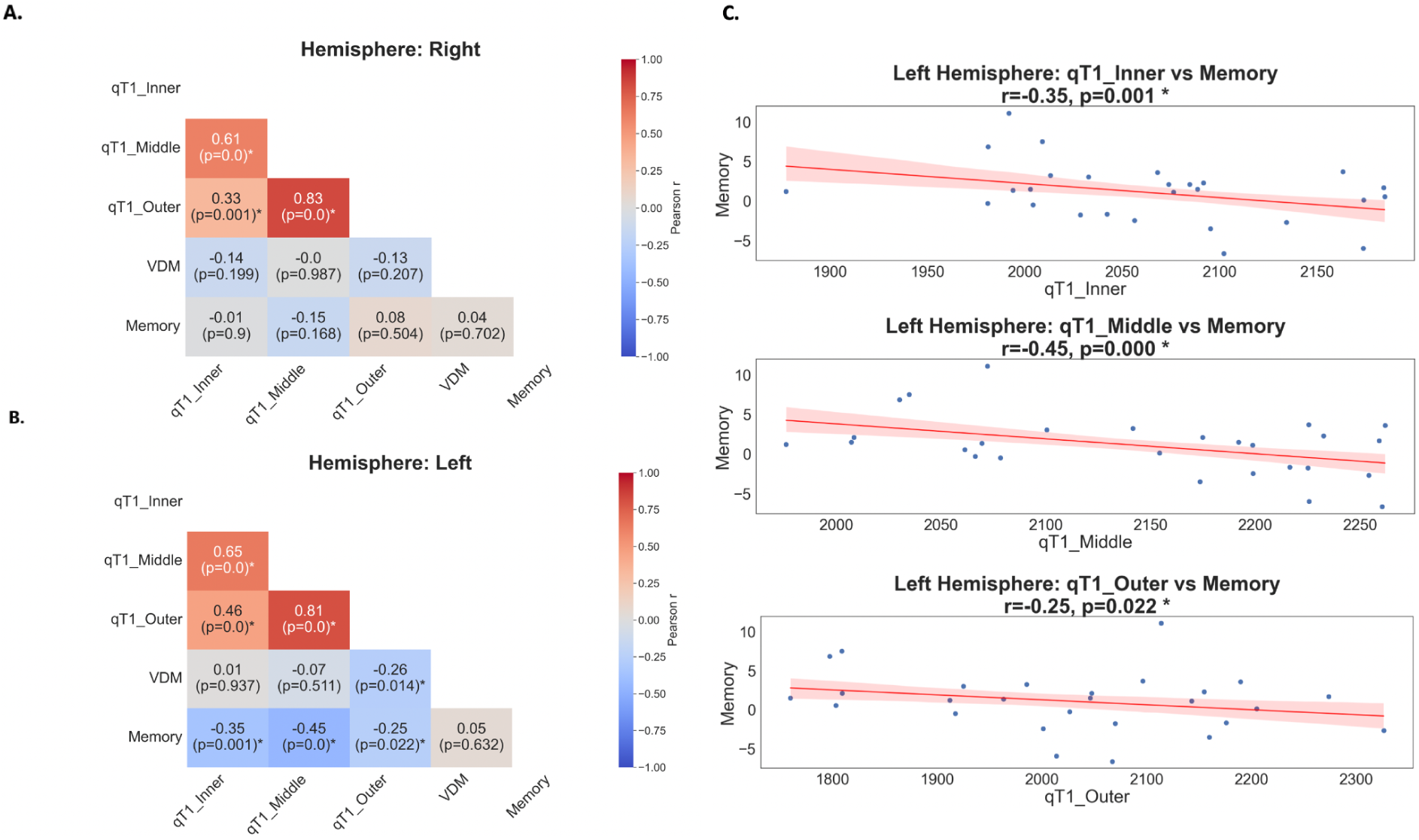
Correlation coefficients and p-values of the relationship between composite memory score and compartment-dependent qT1 values or vessel distance. **A)** Right Hemisphere. **B)** Left Hemisphere. Correlations that survived FDR-correction are marked with an *. **qT1_Inner-Outer** = Mean qT1 times in ms, for compartments 1-3. **VDM** = Vessel Distance in mm. **Memory =** Composite memory score. **C)** Scatter Plots of compartments that showed significant correlations with memory in the left hemisphere.

In order to understand which specific test of the composite memory score drove this effect, we conducted an exploratory analysis and plotted the relationship between the different cognitive tests that formed the composite score and CA1 myelination in different layers (see Supplemental **Figure 6.**). This analysis revealed that the significant correlations with the three layer compartments were mainly driven by the accuracy of the first OLT retrieval (inner × OLT_ret1: *r_FDR_ = -0.43, p < 0.001,* middle × OLT_ret1: *r_FDR_ = -0.43, p < 0.001,* outer × OLT_ret1: *r_FDR_ = -0.29, p = 0.001*), whereas the correlation with qT1 values in the middle compartment was mainly driven by the accuracy in the second OLT retrieval (middle × OLT_ret2: *r_FDR_ = -0.26, p = 0.02*).

We then performed regression analyses to see if myelination in different layers could predict performance in the specific cognitive scores. This analysis revealed a significant effect of memory scores on myelin content *(qT1: β = -9.840 [95% CI -18.033, -1.648], SE = 4.180, p = 0.019)* as well an interaction of this effect with hemisphere (*hemisphere × qT1 interactions: right: β = 9.531 [95% CI 1.734, 17.328], SE = 5.630, p = 0.017).* There was also a trend of higher vessel distance predicting lower cognitive scores, although it did not reach statistical significance *(VDM: β = -9.734 [95% CI -20.767, 1.300], SE = 5.630, p = 0.084)*. To further examine the hemispheric effect, an additional model was used to test for the effect of Memory on qT1 in the left hemisphere only (qT1 ∼ Memory + VDM + Age + Sex + (1+ | Subject)). Additionally, Memory *×* Compartment interactions were tested for the left hemisphere (qT1 ∼ Memory * Compartment + VDM + Age + Sex + (1+ | Subject)) on qT1 (e.g., Memory∼ qT1, Memory ∼ qT1 × hemisphere). Memory predicted qT1 significantly in the left but not right hemisphere. No interaction with the individual compartments was detected (see Supplemental **Table 4**, Supplemental **Table 5,** Supplemental **Table 6** for details). Neither age nor sex related significantly to memory scores (*Age: β = 0.884 [95% CI -7.474,9.242], SE =4.264, p = 0.836,Sex: β = -16.823 [95% CI -85.440, 51.793], SE = 35.009, p = 0.631*).

## Discussion

In the current study, we performed an *in vivo* characterization of the microstructural profile of the hippocampal subfield CA1 in younger adults, using ultra-high-field 7T MRI. We provide evidence for depth-dependent differences in myelination across CA1 compartments that can be detected in-vivo, reflecting the expected three-layered pattern of CA1 known from ex vivo histology. Our study evidences lower qT1 values in right compared to left CA1, and no effect of arterial distance on CA1 myelination. Critically, our data indicates that more myelination is CA1 is linked to better cognitive performance, specifically in object localization. We discuss our results in light of recent evidence on the role of CA1 in cognitive and memory performance in humans.

We show that the degree of myelination, quantified via qT1 relaxation times, differs significantly across CA1 depths. In particular, the inner and outer compartments—bordering the dentate gyrus and cerebrospinal fluid, respectively—show significantly higher degrees of myelination compared to the middle compartment. This finding aligns with the known anatomy of CA1 (Insausti and Amaral 2004; Duvernoy 2004), as the inner and outer compartments of CA1 contain more white matter fibers than the middle compartment, due to the presence the white matter rich structures such as the SRLM in the apical layers, as well as output projections CA1 in the stratum oriens and basal pyramidal layer. The fact that we can visualize this pattern in vivo demonstrates the principle feasibility to detect fine-grained microstructural differences within CA1 with a 7T MRI sequence of 0.5 mm isotropic image resolution, providing a methodological avenue for future research.

Whereas we here measured younger, healthy adults, among whom pathological demyelination is not expected, similar protocols could be applied to understand aging and neurodegeneration. Post mortem studies have shown that early tau accumulation originates in the superficial layers of the entorhinal cortex (Braak and Braak 1991, 1992; Stranahan and Mattson 2010) from which the perforant path (the main source of input to the hippocampus) emerges (Moryś et al. 1994). As these transentorhinal tau stages are common in older adults, one might expect to observe reduced myelination in the inner compartments of CA1 (covering the SRLM) in normal aging, and even more pronounced reductions in AD.

T2-weighted imaging at 7T has evidenced reduced thickness of the SRLM (but not CA1 pyramidal layers) in patients with mild AD compared to controls, and in APOE-e4 carriers compared with noncarriers (Kerchner et al. 2014, 2010; Boutet et al. 2014; Adler et al. 2018). These studies, using 0.2mm^2^ (in-plane) T2-weighted MRI, indicate that the hypointense band corresponding to SRLM is about 0.6 mm to 0.9 mm (Kerchner et al. 2014, 2010; Boutet et al. 2014; Adler et al. 2018) thick in non-demented older adults, and covers approximately the inner quarter of CA1. In our study, the lowest qT1 values—indicating the highest myelination—were observed in the inner compartment (depths 2 to 4, see **Figure 2B**), corresponding to the expected location of the SRLM. In studies of older adults, particularly those with abnormal AD biomarkers, one would expect a flattened U-shape, with pronounced qT1 increases in the innermost layer compartments. Such microstructural alterations could contribute to decreased or aberrant functional connectivity between the superficial layers of the entorhinal cortex and the inner compartment of CA1, which could be studied by means of ultra-high resolution (≤1mm resolution) fMRI (Maass et al. 2014; Zhang et al. 2023).

We here observe significant differences in overall myelination levels of the left and right CA1. This hemispheric difference does not moderate the relationship between CA1 depth and myelination levels. Given myelination predicts cognitive performance, this difference likely reflects underlying biological lateralization. However, B1+ inhomogeneities on the MP2RAGE image intensities could also contribute to this hemispheric asymmetry. While the MP2RAGE is inherently less sensitive to B1+ effects than other sequences (Marques et al. 2010), the degree of that insensitivity depends on specific parameter settings of the acquisition (Haast et al. 2021). Future analyses will have to specifically investigate whether this right-left difference can also be detected in ex vivo histological investigations of the same participants.

Our study does not offer evidence for a relationship between distance to the closest segmented artery and myelination levels in any of the CA1 compartments. This finding contrasts with prior work suggesting that vascular proximity is linked to local microstructure. In the hippocampus, higher myelination has been associated with shorter distance to the supplying vasculature (Haast et al. 2023), consistent with the idea that proximity to arteries supports metabolic supply required for myelin maintenance (Philips and Rothstein 2017). More generally, vascular supply has been related to structural and functional outcomes, including evidence that a dual arterial supply in the hippocampus is associated with better cognitive performance in patients, and relates to gray matter integrity (Perosa et al. 2020; Vockert et al. 2021). While prior work on the neocortex has also reported associations between venous distance and myeloarchitecture across different regions (Knoll et al. 2024), the present results do not indicate such a relationship for CA1 with arterial distance, suggesting that arterial proximity may not be a primary determinant of myelination in this region.

The discrepancy between those and our findings may reflect regional differences. Previous work that investigated the relation between perfusion and arterial blood supply in the hippocampus has found that, among the hippocampal subfields, CA1 shows the greatest distance to supplying macrovasculature, comparatively low microvascular density and consequently the lowest amount of perfusion (Haast et al. 2024). This may suggest that the absence of a detectable association between vasculature and myelination in our study is related to the local architecture of the vasculature, rather than a general absence of a relationship between vasculature and microstructure. In addition, particularly when concerning with very small structures such as CA1, the resolution in which blood vessels can be depicted could make a critical difference. Even at the 0.28 mm isotropic resolution achieved by 7T time-of-flight angiography used here, small arteries that may be critical to innervate small hippocampal subfields may be missed. If certain vessels were not modelled in the automated segmentation process, this could impact the results, too. Future studies may examine how vessel distance is related to (layer-specific) myelination in other hippocampal subfields, as well as older adults, where vascular risk factors and related pathological changes in the brain are usually more prominent (Wahl and Clayton 2024).

Another main motivation of our study was to test for a potential relationship between individual differences in layer-specific myelination in younger adults and individual differences in cognitive performance, specifically memory performance. Pairwise correlations between the composite memory score and depth-dependent myelination revealed significant associations within the left CA1 that survived FDR-correction, indicating that higher myelination levels in left CA1 are related to better cognitive performance. This effect was primarily driven by participants’ retrieval performance in the object-location task, consistent with the view that CA1 plays a key role in memory retrieval processes (Atucha et al. 2023). A linear mixed model analysis revealed that better cognitive performance predicted higher myelination. This effect was further modulated by hemispheric differences, being primarily driven by the left hemisphere, consistent with the initial correlation analysis. This result has interesting implications for future studies. Whereas here, in younger adults, across layer myelination relates to individual cognitive performance, using a combination of task and structural fMRI, it could be investigated in the future if the degradation of specific layers in the course of aging or pathology has specific effects on short versus long-term object retrieval.

The observed pattern of results is consistent with the hypothesis that higher myelination serves as a marker of intact neural function and thus relates to better cognitive performance (Huang et al. 2025). However, the result needs further confirmation by testing a larger cohort with more diverse microstructural and cognitive profiles.

Several limitations should be considered when interpreting these results. Although qT1 mapping is regarded as a reliable method for estimating myelin content—where shorter T1 relaxation times reflect higher myelination (Stüber et al. 2014)—signal specificity could be further improved by incorporating the quantitative susceptibility mapping (QSM), which would help to disentangle myelin contributions from those of other tissue components such as iron (Langkammer et al. 2012). In addition, other MRI approaches exist to study myelination. Quantitative magnetization transfer (qMT) and magnetization transfer saturation (MTsat) model the macromolecular proton pool dominated by myelin lipids and proteins (van der Weijden et al. 2021). Using the MPM, MTsat as well as other qMRI maps can be acquired in a single, fast sequence (Weiskopf et al. 2013). At 7 T MRI, qMT has been pushed to submillimeter resolution (Oh et al. 2018).

For a detailed analysis of CA1 microstructure, qT1 was selected given qT1 maps have been investigated in in-vivo ex-vivo validation studies (e.g. Stüber et al. 2014; Sprooten et al. 2019; Waehnert et al. 2016; Dinse et al. 2013), and provide expected relationships to layer-dependent myelination (e.g. Liu et al. 2025). In addition, it is provided by vendors directly, making it more practical for routine in vivo studies (Bagnato et al. 2018).

Although an 0.5 mm isotropic structural sequence adds significant detail compared to standard 1 mm sequences, novel approaches using 9.4 or even >10 Tesla allow acquiring MP2RAGE sequences at 0.4 mm image resolution (van der Zwaag et al. 2025). Such non-standard acquisitions could, in a small number of subjects, be used to validate the presented methods. Additionally, we here focused on CA1, where most memory tasks are mediated by a network of brain areas. The correlation between the object localization task and CA1 myelination indicates that for network-level investigations, CA1 should be taken into account in addition to CA3 and the EC. Both the hippocampal head and tail were excluded from analysis in this study. However, the hippocampus exhibits pronounced functional differentiation along its longitudinal (anterior–posterior) axis, with anterior and posterior segments contributing differentially to cognitive and memory processes (see e.g. (Xie et al. 2024; Angeli et al. 2025)). Accordingly, future work aiming to further elucidate microstructural contributions to cognition should also consider microstructural variation of the CA1 subfield along its longitudinal axis (Tang et al. 2020).

Finally, it is worth mentioning that we performed manual co-registration for qT1 to T2 registration, and we also manually checked for the correct segmentation of the hippocampal subfields (see also Canada et al. 2024). While these approaches ensure high quality, there is the need to develop tools that produce robust results without manual intervention to ensure reproducibility.

Taken together, future studies may use the here presented knowledge, methodology and insight of the in-vivo architecture of CA1 described with 7T MRI and its relation to cognitive performance to study depth-dependent myelination in normal aging and AD, where perforant pathway input to apical layers of CA1 might be degraded. Specifically, these studies may determine whether or not tau burden is related to reduced myelination in inner CA1 layers, and whether or not myelination is associated with memory deficits or decline. Furthermore, extending this approach to the entire hippocampus and parahippocampal regions, and integrating it with functional MRI, may enable the investigation of depth-specific functional connectivity patterns and their alterations with aging or disease progression and cognition.

## Acknowledgments

This project has received funding from the European Research Council (ERC) under the European Union’s Horizon 2020 research and innovation programme (grant greement No 949609).

## Footnotes

1 The code used for study is available at: https://github.com/Zeconyser/The-In-vivo-Microstructural-Profile-of-Human-Hippocampal-Subfield-CA1/tree/main

