## Supplementary Material for "The in-vivo microstructural profile of human hippocampal subfield CA1 and its relation to memory performance"

**SUPPLEMENTARY**

### Supplementary figures

#### MRI Protocol MTLVESSEL

| Sequence | MP2RAGE | T2w TSE | ToF | RS - fMRI | CVR - fMRI |
|---|---|---|---|---|---|
| Resolution (mm) | 0.5 x 0.5 x0.5 | 0.44 x 0.44 x 1.1 | 0.28 x 0.28 x 0.28 | 0.9 x 0.9 x 0.9 | 0.9 x 0.9 x 0.9 |
| FoV read (mm) x FoV phase (%) | 240.0 x 100.0 | 224.0 x 100.0 | 200.0 x 87.5 | 212.0 x 99.2 | 212.0 x 100.0 |
| TR (ms) | 4800.0 | 8000.0 | 22.0 | 2000.0 | 2000.0 |
| TE (ms) | 2180.0 | 92.0 | 4.59 | 20.0 | 20.0 |
| Number of slices | 352 | 50 | 176 | 58 | 58 |
| TI 1/TI 2 (ms) | 900.0 / 2750.0 | N.A. | N.A. | N.A. | N.A. |
| Flip angle (°) | 5.0 / 3.0 | 60.0 | 19.0 | 80.0 | 80 |
| Bandwidth (Hz/px) | 250.0 | 158.0 | 142.0 | 1412.0 | 1412.0 |
| GRAPPA | 2 | N.A. | 3 | 4 | 4 |
| Length (min) | 17:14 | 07:38 | 14:03 | 05:03 | 08:55 |

***Figure 1.*** *FoV = Field of View, TR = Repetition time, TE = Echo time, TI 1/ TI 2 = T1/T2 Relaxation times,*
*GRAPPA = Generalized autocalibrating partially parallel acquisitions (acquisition acceleration method), add abbrevs for sequences*

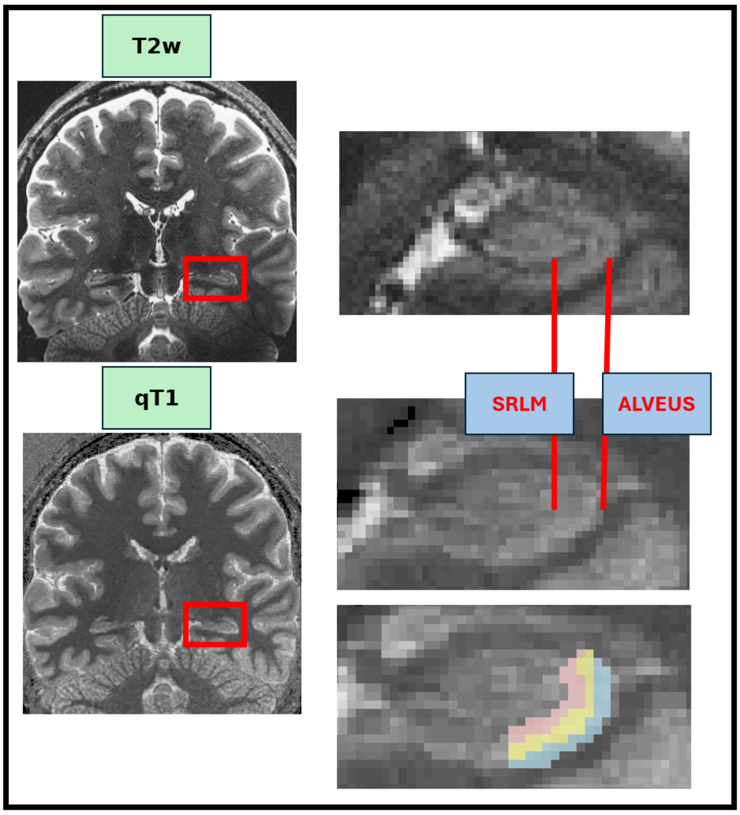


***Figure 2.*** *Comparison of Hippocampus contrasts on qT1 and T2w scan. The laminar boundaries of Subfield CA1 (Alveus towards the outer compartment, and SRLM at inner compartment) are indicated. CA1 Mask, derived from ASHS segmentation, covers the subfield between these boundaries.*

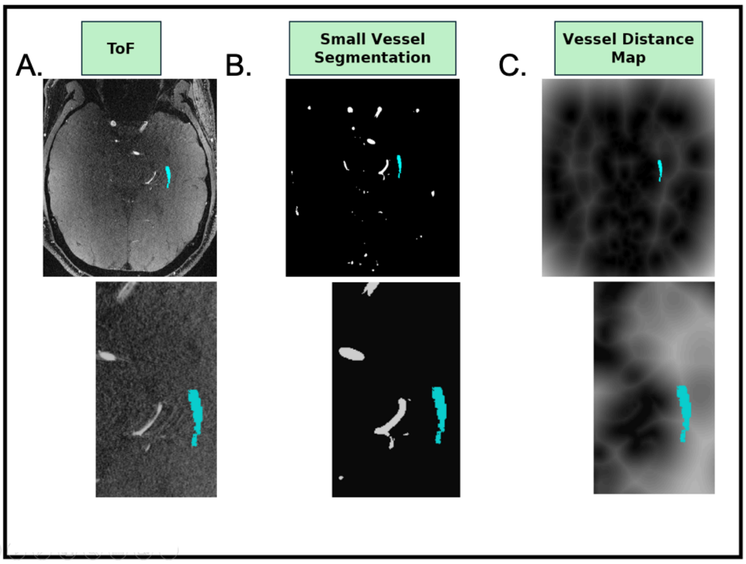


***Figure 3. Detailed overview over vessel distance analyses.*** *Coronal views (images) of each step in the vessel distance analysis. Binary mask of CA1 (head and tail excluded) shown in blue for visualization.* ***A)*** *Time of Flight Angiography (ToF)* ***B)*** *Binary vessel segmentation mask obtained from the OMLETTE toolbox, with $\tau$ = 0.5* ***C)*** *Global vessel distance map, in which each voxel carries the information about the euclidean distance to the nearest segmented vessels in mm.*

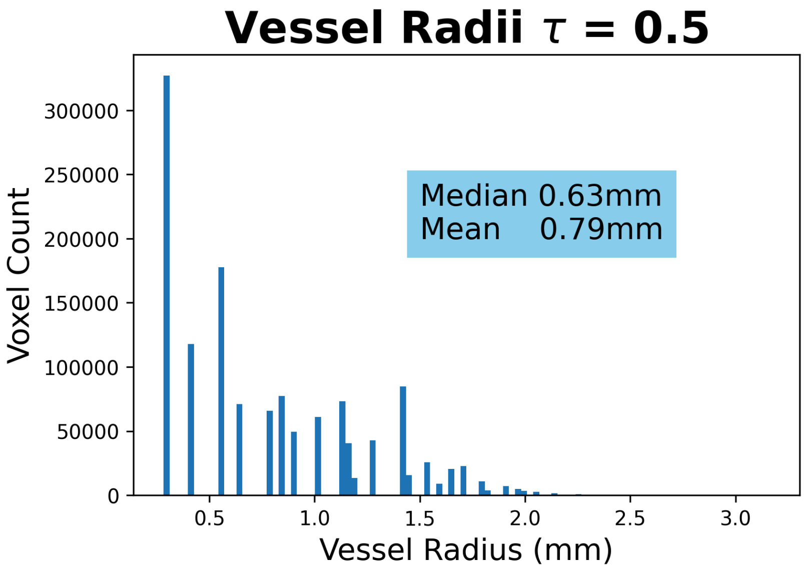


***Figure 4.*** *Distribution of radii of segmented vessels for the whole slab, across all participants.. Voxel count refers to the number of vessel skeleton voxels that have a specific radius to the vessel border.*

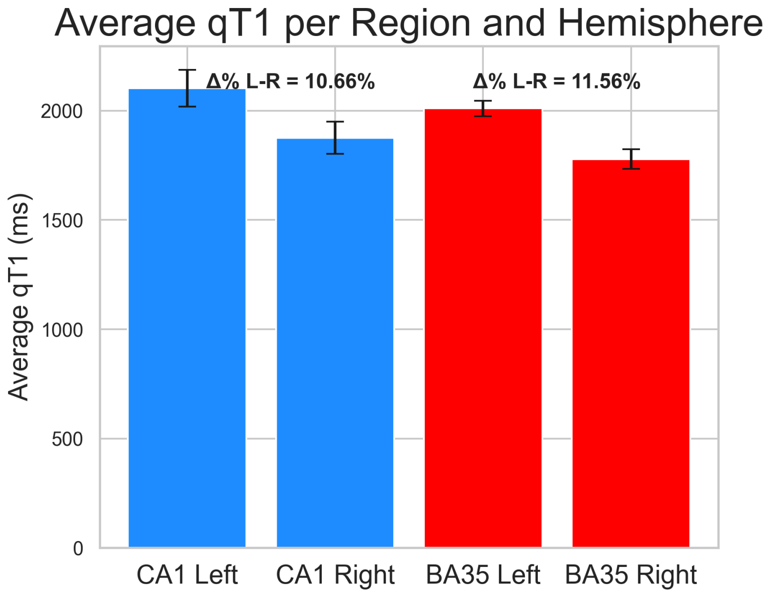


***Figure 5.*** *Differences in average qT1 (ms) between hemispheres for CA1 and BA35 as a control region. Both regions demonstrate lower average qT1 (implying higher myelination) in the right hemisphere compared to the left.*

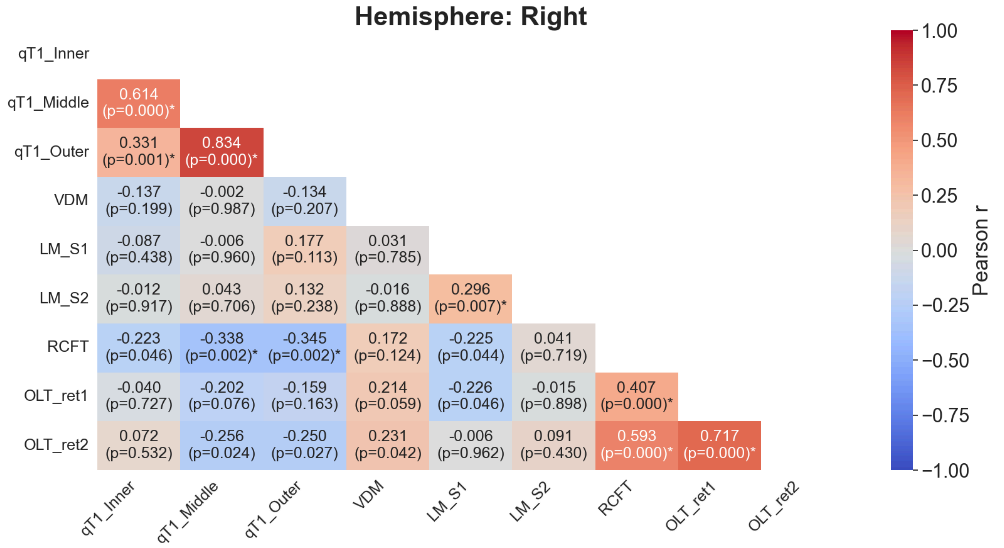

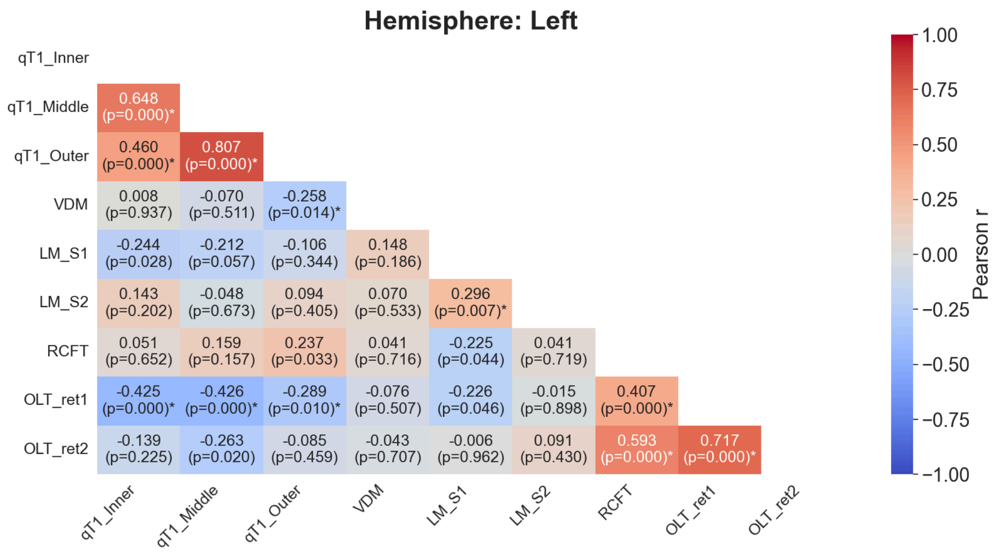


***Figure 6**. Raw correlations and p-values of individual cognitive test scores with compartment-depent qT1 values and Vessel distance. Correlations that were still statistically significant after being FDR corrected are marked with an (*). **qT1_Inner-Outer** = Mean qT1 times in ms, for the three layer compartments. **VDM** = Vessel Distance in mm. **LM_S1/S2** = Total Logical Memory score over both recalls (immediate and delayed), for both stories (S1/S2). **RCFT** = Total Rey Complex Figure Task score for recall. **OLT_ret1/ret2** = Total Object Location Task accuracy in both retrievals.*

### Supplementary tables

#### Table 1. Linear mixed model (qT1 ~ Compartment * Hemisphere + Age + Sex)

| *Predictors* | *Estimates* | *std. Error* | *CI* | *z* | *p* |
|---|---|---|---|---|---|
| (Intercept) | 2031.41 | 108.11 | 1819.51 –2234.31 | 18.78 | <0.001 |
| Compartment [Middle] | 82.00 | 24.46 | 32.09 – 131.90 | 3.22 | 0.001 |
| Compartment [Outer] | -24.49 | 25.46 | –74.40 – 25.41 | -0.96 | 0.336 |
| Hemisphere [Right] | -226.72 | 25.46 | -276.63 –176.82 | -8.90 | <0.001 |
| Sex [Male] | -2.749 | 34.68 | -70.73 – 65.23 | -0.08 | 0.94 |
| Compartment[Middle]/Hemisphere[Right] | -31.57 | 36.01 | -102.15 – 39.00 | -0.87 | 0.38 |
| Compartment[Outer]/Hemisphere[Right] | 24.89 | 36.01 | -45.67 – 95.47 | 0.69 | 0.49 |
| Age | 1.48 | 4.36 | -7.07 – 10.03 | 0.34 | 0.73 |
| Group Variance | 3462.14 | 18.11 | | | |
| Observations | 144 | | | | |

#### Table 2. Linear mixed model (qT1 ~ Compartment * VDM + Age + Sex)

| *Predictors* | *Estimates* | *std. Error* | *CI* | *z* | *p* |
|---|---|---|---|---|---|
| (Intercept) | 2064.14 | 12.46 | 1814.74 –2313.54 | 16.22 | <0.001 |
| Compartment [Middle] | 73.76 | 80.08 | -83.19 – 230.72 | 0.92 | 0.36 |
| Compartment [Outer] | 115.43 | 82.14 | –45.56 – 276.42 | 1.40 | 0.16 |
| Hemisphere [Right] | -226.87 | 14.70 | -255.69 – -198.05 | -15.43 | <0.001 |
| Sex [Male] | -2.81 | 34.50 | -70.44 – 64.82 | -0.08 | 0.93 |
| VDM | -2.08 | 8.21 | -18.16-14.00 | -025 | 0.80 |
| VDM/Compartment[Middle] | -0.92 | 10.36 | -21.23 – 19.39 | -0.09 | 0.93 |
| VDM/Compartment[Outer] | -15.52 | 10.32 | -35.75 – 4.71 | -1.50 | 0.13 |
| Age | 0.82 | 4.36 | -7.24 – 9.37 | 0.19 | 0.85 |
| Group Variance | 3423.08 | 18.00 | | | |
| Observations | 144 | | | | |

#### Table 3. Linear mixed model (qT1 ~ Memory * Hemisphere + VDM + Age + Sex)

| *Predictors* | *Estimates* | *std. Error* | *CI* | *z* | *p* |
|---|---|---|---|---|---|
| (Intercept) | 2158.383 | 117.913 | 1927.278 –2389.488 | 18.305 | 0.000 |
| Hemisphere [Right] | -237.816 | 16.284 | -269.733 – -205.900 | -14.604 | <0.01 |
| Sex [Male] | -16.823 | 35.009 | -85.440 – 51.793 | -0.481 | 0.631 |
| Memory | -9.840 | 4.180 | -18.033 – -1.648 | -2.396 | 0.019 |
| Memory/Hemisphere[Right] | 9.531 | 3.978 | 1.734-17.328 | 2.396 | 0.017 |
| VDM | -9.734 | 5.630 | -20.767 - 1.300 | -1.729 | 0.084 |
| Age | 0.884 | 4.264 | -7.474 – 9.242 | 0.207 | 0.863 |
| Group Variance | 3046.396 | 16.507 | | | |
| Observations | 144 | | | | |

#### Table 4. Linear mixed model (qT1 ~ Memory + VDM + Age + Sex) - Left Hemisphere

| *Predictors* | *Estimates* | *std. Error* | *CI* | *z* | *p* |
|---|---|---|---|---|---|
| (Intercept) | 2170.894 | 183.874 | 1810.508 - 2531.279 | 11.806 | 0.000 |
| Sex [Male] | -35.968 | 47.295 | -128.664 – 56.729 | -0.760 | 0.447 |
| Memory | -10.468 | 4.913 | -20.097 – -0.0839 | -2.131 | 0.033 |
| VDM | -12.369 | 10.674 | -33.290 – 8.553 | -1.159 | 0.247 |
| Age | 1.533 | 5.893 | -10.018 - 13.084 | 0.260 | 0.795 |
| Group Variance | 5213.638 | 33.666 | | | |
| Observations | 72 | | | | |

#### Table 5. Linear mixed model (qT1 ~ Memory + VDM + Age + Sex) - Right Hemisphere

| *Predictors* | *Estimates* | *std. Error* | *CI* | *z* | *p* |
|---|---|---|---|---|---|
| (Intercept) | 1977.565 | 165.259 | 1653.663 - 2301.467 | 11.996 | 0.000 |
| Sex [Male] | 11.363 | 49.869 | -86.360 - 109.087 | 0.228 | 0.820 |
| Memory | 0.736 | 4.850 | -8.770 - 10.243 | 0.152 | 0.897 |
| VDM | -13.033 | 10.737 | -34.078 - 8.012 | -1.214 | 0.225 |
| Age | -0.724 | 5.636 | -11.770 - 10.322 | 5.636 | -0.128 |
| Group Variance | 5746.866 | 40.784 | | | |
| Observations | 72 | | | | |

**Table 6. Linear mixed model (qT1 ~ Memory * Compartment + VDM + Age + Sex) - Left Hemisphere**

| *Predictors* | *Estimates* | *std. Error* | *CI* | *z* | *p* |
|---|---|---|---|---|---|
| (Intercept) | 2099.719 | 183.889 | -1739.304 –2460.134 | 11.418 | 0.000 |
| Compartment[Middle] | 87.975 | 23.287 | 42.333 - 133.618 | -0.00 | 1.00 |
| Compartment[Outer] | -14.612 | 24.377 | -62.390 - 33.167 | -0.599 | 0.549 |
| Sex [Male] | -38.744 | 47.230 | -131.313 – 53.825 | -0.820 | 0.412 |
| Memory | -8.272 | 5.891 | -19.818 - 3.274 | -1.404 | 0.160 |
| Memory/Compartment[Middle] | -3.285 | 5.656 | -14.371 - 7.800 | -0.581 | 0.561 |
| Memory/Compartment[Outer] | -3.424 | 5.656 | -14.510 - 7.661 | -0.605 | 0.545 |
| VDM | -8.233 | 10.946 | -29.688 – 13.221 | -0.752 | 0.452 |
| Age | 2.168 | 5.894 | -9.385 – 13.721 | 0.368 | 0.713 |
| Group Variance | 6023.335 | 40.365 | | | |
| Observations | 72 | | | | |
